# Unbiased classification of mosquito blood cells by single-cell genomics and high-content imaging

**DOI:** 10.1101/234492

**Authors:** Maiara S. Severo, Jonathan J.M. Landry, Randall L. Lindquist, Christian Goosmann, Volker Brinkmann, Paul Collier, Anja E. Hauser, Vladimir Benes, Johan Henriksson, Sarah A. Teichmann, Elena A. Levashina

**Affiliations:** Vector Biology Unit, Max-Planck-Institute for Infection Biology, Charitéplatz 1, 10117 Berlin, Germany; Genomics Core Facility, European Molecular Biology Laboratories, Meyerhofstraße 1, 69117 Heidelberg, Germany; Deutsches Rheumaforschungszentrum, Charitéplatz 1, 10117 Berlin, Germany; Microscopy Core Facility, Max-Planck-Institute for Infection Biology, Charitéplatz 1, 10117 Berlin, Germany; Immune Dynamics and Intravital Microscopy, Charité Universitätsmedizin, Charitéplatz 1, 10117 Berlin, Germany; Wellcome Trust Sanger Institute and European Bioinformatics Institute, Wellcome Genome Campus, Hinxton, Cambridge, CB10 1SD, UK; Lead contact

## Abstract

Mosquito blood cells are ancestral immune cells that help control infection by vector-borne pathogens. Despite their importance, little is known about mosquito blood cell biology beyond the ambiguous morphological and functional criteria used for their classification. Here we combined the power of single-cell RNA-sequencing, imaging flow cytometry and single-molecule RNA hybridization to analyze blood cells of the malaria mosquito *Anopheles gambiae*. By demonstrating that blood cells express nearly half of the mosquito transcriptome, our dataset represents an unprecedented view into their transcriptional machinery. Analyses of differentially expressed genes identified transcriptional signatures of two distinct cell types that challenge the current morphology-based classification of these cells. We further demonstrated an active transfer of a cellular marker between blood cells that confounds their identity. We propose that cell-to-cell exchange is broadly relevant for cell type classification and may account for the remarkable cellular diversity observed in nature.

## INTRODUCTION

The cell is the basic building block of all living organisms. All the major decisions coordinating systems at the organismal level, be it life or death, health or disease, start within a cell. In eukaryotes, multicellularity came with cell compartmentalization and specialization. Cells found in the blood, or hemolymph in invertebrates, have received numerous denominations, such as hemocytes, amebocytes, phagocytes, coelomocytes and immunocytes (Ottaviani and Franceschi, 1997). Regardless of their names, these cells play key roles in shaping the extracellular environment and helping fight infection all throughout the animal kingdom. In insects, blood cells are found circulating by the flow of hemolymph, or as sessile cells associated with internal organs (Ribeiro and Brehelin, 2006). These cells are considered the equivalent of leukocytes in mammals, and display extraordinary functional resemblances to neutrophils, monocytes and macrophages, e.g. phagocytic abilities, chemotaxis, production of antimicrobial peptides, free radicals and cytokine-like molecules (Bergin et al., 2005; Browne et al., 2013; Buchmann, 2014; Costa et al., 2005; Lavine and Strand, 2002). Contrary to the well-established classification of human leukocytes, blood cell type classification is controversial in insects, with similar terms being used for different cell morphologies even within the same insect order (Brayner et al., 2005; Castillo et al., 2006; Hernandez et al., 1999; Ribeiro and Brehelin, 2006). Of note, most studies of insect blood cells have focused on the embryonic and larval stages of the *Drosophila* model (Brandt et al., 2008; Vlisidou and Wood, 2015; Wood and Jacinto, 2007) and these observations are not immediately applicable to other insects (Ribeiro and Brehelin, 2006; Zdobnov et al., 2002), particularly in the study of the adult life stages of insects implicated in disease transmission to humans.

Mosquitoes are the deadliest animals on Earth, transmitting pathogens that cause a variety of diseases and infect millions of people worldwide every year (WHO, 2014). While feeding on blood to reproduce, adult female mosquitoes acquire blood-borne pathogens from an infected host. Pathogen development and replication within the mosquito is an absolute requirement for further transmission, so the disease cycle is, in part, determined by the mosquitoes’ capacity to counterattack these pathogens. Mosquito blood cells are of vital importance in this process as they represent the cellular arm of mosquito immunity, and also participate in humoral responses by secreting pathogen-killing factors, such as components of the melanization pathway (Hillyer et al., 2003; Yassine et al., 2012) and of the complement-like system that help eliminate malaria parasites (Blandin et al., 2004; Frolet et al., 2006). Earlier transcriptomics studies based on microarrays have explored the molecular basis of mosquito hemocyte immunity upon infection with bacteria and *Plasmodium* (Baton et al., 2009; Pinto et al., 2009). More recently, Smith et al used mass spectrometry to analyze mosquito hemocytes isolated based on their uptake of magnetic beads (Smith et al., 2016). To identify genes and proteins more predominantly expressed in hemocytes or in ‘phagocytes’, these studies relied on enrichment analyses of blood cells relative to whole body or total hemolymph samples, respectively. This restricted the identification of genes co-expressed in blood cells and other tissues, and masked the contribution of sessile hemocytes to overall systemic responses. Such approaches are standard in the study of hemocyte biology, mainly due to the practical constraints associated with the scarcity of these cells and the lack of cellular markers for successful hemocyte isolation and purification.

Importantly, the absence of cellular markers has also precluded the classification of mosquito blood cells beyond morphology and function. To date, mosquito hemocytes are morphologically divided based on label-free light microscopy into: (1) granulocytes, phagocytic cells that exhibit granules in their cytoplasm and quickly spread onto glass; (2) oenocytoids, spherical and poorly adhesive cells that produce phenoloxidases involved in melanization defenses; and (3) prohemocytes, small cells of reportedly 2 μm (Rodrigues et al., 2010; Smith et al., 2015) or 4–6μm in size that have been suggested to function as hemocyte progenitors (Castillo et al., 2006), and/or represent small phagocytic cells arising from asymmetrical divisions (King and Hillyer, 2013). Although it has been demonstrated that mosquito hemocytes increase in numbers upon blood feeding and infection, in the absence of known hematopoietic organs (Bryant and Michel, 2014; King and Hillyer, 2013), the pathways underlining their differentiation into these three classes remain unknown. Whether the current classification represents true discrete cell types or states, and if mosquito blood cell subpopulations exist, are also yet to be explored. In humans and mice, single-cell transcriptomics have recently began to tackle similar questions. It is now also increasingly evident that significant functional differences and considerable variability in gene expression can be recognized in cells long considered to be of the same type (Gaublomme et al., 2015; Grun et al., 2015; Shalek et al., 2014). The use of single-cell approaches to explore cellular heterogeneity in non-model organisms holds the promise to uncover unforeseen complexity, and identify cellular populations that would be forever ‘masked’ in bulk, enrichment-based measurements. In addition, single-cell studies of invertebrates can contribute to comparative analyses of cellular diversity across different systems.

Here we unravel the molecular fingerprint of a subset of mosquito blood cells at a single-cell level, and show that naïve, unstimulated hemocytes express nearly half of the mosquito transcriptome. Our dataset represents a valuable resource for further studies of transcriptional regulation in mosquitoes, and paves the way for the identification of new molecules controlling infection by vector-borne pathogens. By applying fluorescence-activated cell sorting (FACS), single-cell RNA-sequencing (scRNA-seq) and high-content imaging flow cytometry to the analysis of mosquito hemocytes, our unparalleled study identifies two distinct blood cell subpopulations, which we propose to call plasmatocytes and melanocytes. Remarkably, the identified cell subsets do not directly correspond to the current morphological and functional classification of mosquito blood cells, and indicate that the emerging observations of cellular heterogeneity in humans and mice are also characteristic of ancestral blood cells. Our findings further reveal active molecular exchange between mosquito blood cells and the presence of extracellular vesicles (EVs) in the mosquito hemolymph. Shuffling of mRNA and proteins between distinct cell types can influence cellular identity and, in turn, confound cell type identification and classification. Altogether, we demonstrate the power of single-cell technologies in shedding light on the contribution of cell-to-cell exchange to cellular diversity and provide a new perspective on the discussion of the slippery concept of ‘cell types’.

## RESULTS

### Single-cell RNA-sequencing of blood cells from *PPO6::RFP* transgenic mosquitoes

In the absence of hemocyte-specific antibodies and dyes, or of transgenic mosquitoes expressing pan-hemocyte markers, we chose to explore a subset of blood cells identified in a transgenic mosquito strain expressing a red fluorescence reporter (tdTomato, abbreviated herein as RFP) under the control of the *prophenoloxidase 6 (PPO6)* melanization-related gene (*PPO6::RFP*) (Volohonsky et al., 2015). Melanization is a well-established immune response of invertebrates that controls infection against bacteria and parasites (Abraham et al., 2005; Christensen et al., 2005; Hillyer and Strand, 2014; Michel et al., 2005). Several reports suggest that melanization is mediated by a specific cell population called oenocytoids, which represents approximately 10% of the blood cells (Castillo et al., 2006; Hillyer and Strand, 2014), but these cells have not been directly explored. Our focus was on cells obtained in the absence of infection or blood feeding, i.e. during homeostasis because it provides a baseline for analysis of cell-to-cell variation. Using this transgenic strain, we first investigated whether RFP-positive hemocytes were present in the mosquito circulation. For that, we perfused hemolymph onto microscope slides and identified cells displaying RFP fluorescence within the size range predictive of hemocytes (Figure 1A). RFP signal was also observed in hemocytes attached to the inner abdominal wall of dissected adult female mosquitoes, where fat body cells are most prominent (Figure 1B, arrow). As expected, the hemolymph perfusate was, however, heavily contaminated with a mixture of cells and subcellular/tissue debris (Castillo et al., 2006). To purify live RFP-positive cells, we developed a FACS approach based on RFP expression and Hoechst nuclear staining, and validated our purification method by microscopic inspection of sorted cells (Figure 1C-D). Notably, the sorted cell population corresponded to 0.1% of the total events measured (n=100,000) in the perfusate of at least 10 mosquitoes (see Methods). This is in accordance with previous work indicating that only a small subset of adult mosquito hemocytes produce PPOs (Hillyer and Strand, 2014).

**Figure 1:**
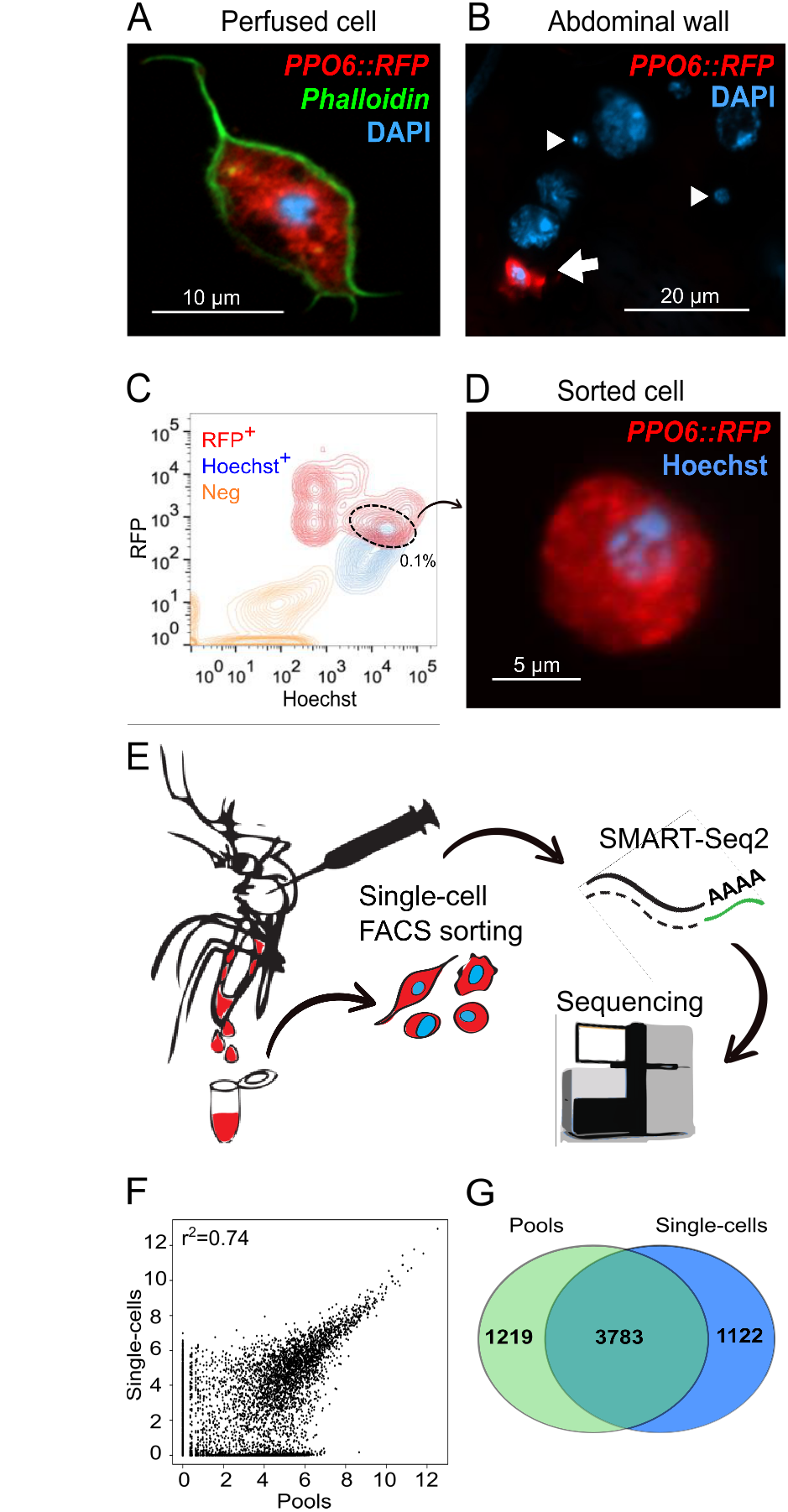
Single-cell RNA-sequencing of transgenic mosquitoes expressing a RFP reporter. A *PPO6::RFP* transgenic mosquito strain was used for isolation of blood cells. (A) A RFP-expressing hemocyte (red) obtained by perfusion of a female adult mosquito. (B) Only a proportion of adult mosquito blood cells display RFP expression (red, arrow), whereas other cells of sizes suggestive of hemocytes do not (arrowhead). (C) FACS sorting of RFP^+^ (R^+^) Hoechs^+^ (H^+^) blood cells from hemolymph of perfused *PPO6::RFP* mosquito females. (D) Representative image of a sorted cell. (E) The pipeline developed for the study of mosquito blood cells based on single-cell RNA-sequencing. Cells were sorted into a 96-well plate, processed according to the SMART-Seq2 protocol and sequenced in a HiSeq Illumina platform. (F) Scatter plot for the average normalized read counts from pools and single-cells. r^2^ indicates Pearson correlation. (G) Venn diagram of genes detected in single-cells and pools (normalized count ≥ 1). Scale bars: (A) 10 μm, (B) 20 μm, (D) 5 μm. DNA is stained with DAPI (A-B) and Hoechst (D).

Next, we FACS-sorted single blood cells and performed scRNA-seq to capture the transcriptome of single PPO-producing blood cells (Figure 1E). Hemocytes were sorted into a 96-well plate and after sample processing and quality assessment (see Methods), we obtained successful cDNA amplification for 56 single-cells in addition to two pools of 30 cells each, from which 28 high quality cDNA libraries were sequenced, representing 26 single hemocytes and the two pooled samples (Figure S1A-D). As a single mosquito can contain as little as 500 blood cells in the circulation (Bryant and Michel, 2014; Hillyer, 2010), we believe the small number of cells analyzed reflects a combination of technical limitations inherent to our approach. First, blood cell isolation is not trivial in mosquitoes, as it requires hemolymph perfusion under micromanipulation followed by cell purification, posing great difficulty in obtaining high numbers of live cells. Second, mosquito blood cells reportedly vary in size from as little as 2 to 20 μm (Brayner et al., 2005; Bryant and Michel, 2016; Castillo et al., 2006; Hernandez et al., 1999), and variability in cell size can significantly affect RNA recovery, as small cells contain small amounts of RNA. The adaptation of the scRNA-seq protocol to the study of an invertebrate system, e.g. the choice of lysis buffer and chemistry, may also have influenced our results, especially since different biological cell types show distinct technical quality features in scRNA-seq experiments (Ilicic et al., 2016). A small number of cells has, nevertheless, been used in other scRNA-seq studies (Shalek et al., 2013; Xue et al., 2013) and does not preclude identification of cellular types when an adequate sequencing depth is used. We, therefore, prioritized the deep sequencing of individually curated, very high-quality samples representing a small subset of cells obtained *ex vivo*.

Indeed, our sequencing generated on average 4.5 million reads per sample, well-above the minimum of one million reads previously suggested as a requirement for adequate single-cell studies (Wu et al., 2014). Over 70% of the reads were successfully mapped, with exonic reads comprising of more than 40% (Figure S1E-F and Table S1). All samples achieved saturation at around 2 million reads, comparable to previously observed for mammalian cells (Wu et al., 2014). For further analyses, we discarded one cell as it showed gene expression suggestive of a doublet (Figure S1F). Doublets have been reported in the circulation of adult female mosquitoes (King and Hillyer, 2013) and may fall within a size range comparable to larger hemocytes. Around 3,800 genes were detected in each pool, whereas single-cells expressed between 450 and 1,400 genes. Similar expression profiles were obtained for pools and single-cells (Figure 1F) with comparable numbers of detected genes only in single-cells or in the pools (1,100 and 1,200 genes, respectively) (Figure 1G). The marker genes used for FACS-sorting (*PPO6* and *tdTomato*) were identified in both single-cells and pools (Table S2), confirming the efficiency of our method. Altogether, our results show that the transcriptome of mosquito hemocytes comprises of over 6,000 genes, of which more than half (3,400) had not been identified in earlier studies (Figure S1G) (Baton et al., 2009; Pinto et al., 2009). In addition, our approach revealed sequences for over 80% (n=914) of the proteins reported by an earlier proteomics approach based on magnetic beads isolation of *A. gambiae* phagocytes (Smith et al., 2016) (Figure S1H), corroborating our findings and demonstrating the higher sensitivity of RNA sequencing as compared to proteomics.

Mosquito hemocyte biology has been mostly studied in the context of immunity. We, thus, inspected our dataset for previously identified immune genes. Several members of immune pathways were expressed at low levels in some naïve hemocytes, such as the transcription factors *REL1* (AGAP009515) and *REL2* (AGAP006747), *Cactus* (AGAP007938), *I_ĸ_B_γ_* (AGAP009166) and *I_ĸ_B_γ_* (AGAP005933), and the receptors *PGRP-LC* (AGAP005203) and *PGRP-S1* (AGAP000536) (Table S2). Components of the complement cascade, e.g. *TEP1* (AGAP010815), *APL1C* (AGAP007033), *LRIM1* (AGAP006348) and *HPX2* (AGAP009033), were also detected in some cells, along with the LPS-induced TNFα transcription factor (LITAF)-like 3 (AGAP009053) described to control *Plasmodium* survival in the gut (Smith et al., 2015). The phagocytic and antibacterial activities of these cells can be illustrated by the expression of *Eater* (AGAP012386), *Ninjurin* (AGAP006745) and *Nimrod* (AGAP009762), alongside that of several fibrinogen-related proteins (FREPs/FBNs), such as *FBN8* (AGAP011223), *FBN9* (AGAP011197), *FBN10* (AGAP011230) and *FBN30* (AGAP006914) (Dong and Dimopoulos, 2009; Estevez-Lao and Hillyer, 2014; Lombardo et al., 2013). Although no ortholog for a major *Drosophila* hemocyte marker, hemolectin, has been described in the *A. gambiae* genome, mosquito hemocytes expressed both *Pannier* (AGAP002235) and *Serpent* (AGAP002238) GATA factors, as well as *misshapen* (AGAP006340), which are associated with blood cell differentiation, maturation and activation in fruit fly larvae (Braun et al., 1997; Fossett et al., 2003; Minakhina et al., 2011). Genes involved in cell adhesion and polarity, such as *integrin β−1* (AGAP010233), *laminin* (AGAP010548), *Notch1* (AGAP001015) and *Armadillo* (AGAP001043), and components of extracellular matrix like *collagen type IV*(AGAP009200) were also identified. The processed gene expression data for visualization in single cells is accessible at http://data.teichlab.org. Our transcriptional data suggests that in addition to immunity, naïve blood cells are performing tissue maintenance and morphogenesis tasks.

### Identification of blood cell subpopulations

To account for the technical noise arising from the small amounts of RNA, we included in our samples External RNA Controls Consortium spike-ins (ERCC) prior to cDNA amplification (Baker et al., 2005). We analyzed percentage of spike-ins and mitochondrial counts as a proxy for sequencing efficiency, RNA degradation or incomplete lysis and potential cell death. As anticipated, variation was observed (Figure S1I-J), but caution was taken in applying these criteria and attributing them biological meaning because variability could have arisen from true cell type-related processes. Differences in total number of expressed genes could also have stemmed from different morphologies and cell types. Therefore, we decided not to further discard any cells, and manually curated their individual mappings to confirm that the samples corresponded to potentially true representations of mosquito blood cells. To estimate technical noise, we applied the variability threshold based on the square of the coefficient of variation (CV^2^) of the spike-ins (Brennecke et al., 2013), and identified 148 genes whose expression exceeded the variability threshold modeled by the spike-ins (Figure 2A). These highly variable genes included a scavenger receptor, fibrinogen-related and leucine-rich repeat-containing proteins, as well as genes involved in vesicle transport, metabolism and transcription (Table S3). No genes directly associated with cell cycle had high variability, although several *cyclin* genes were detected in specific cells (Table S2), corroborating previous reports of the potential of mosquito hemocytes to undergo cellular division (Bryant and Michel, 2014, 2016; King and Hillyer, 2013).

**Figure 2:**
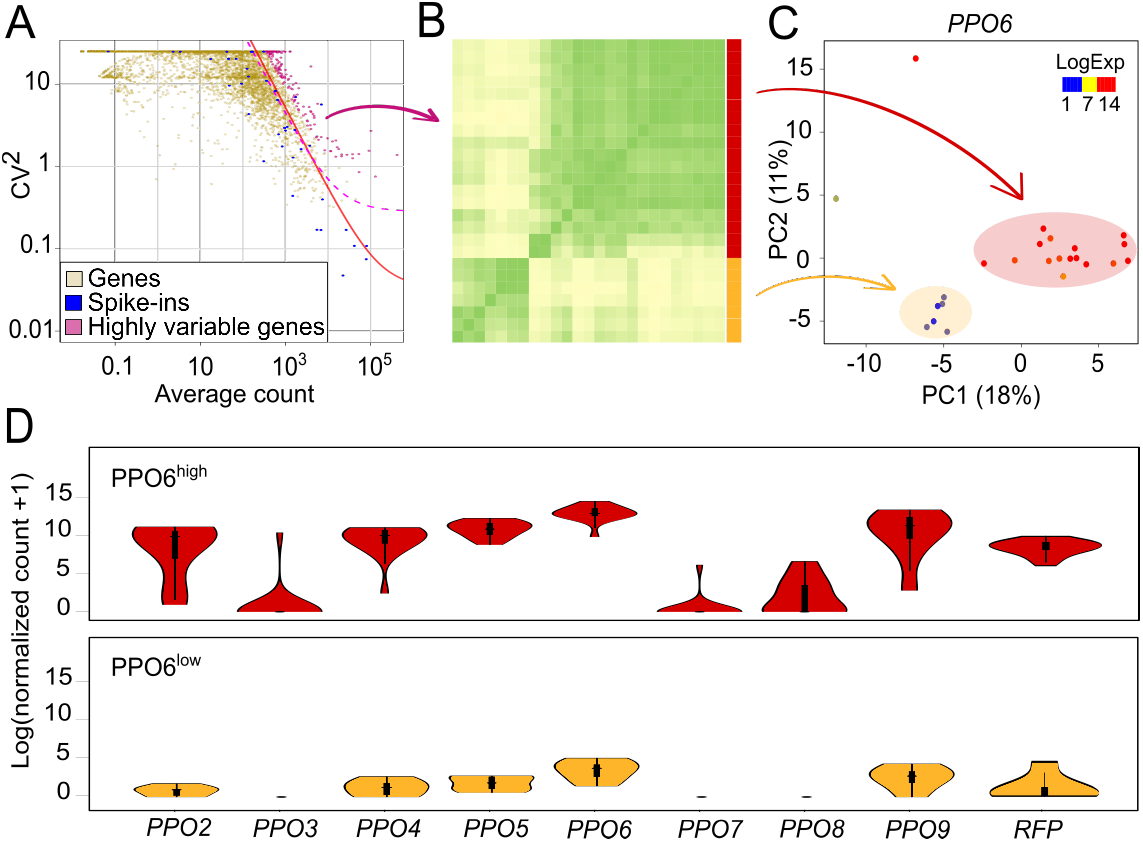
Identification of mosquito blood cell subpopulations. (A) The expression variability of individual genes measured by the squared coefficient of variation (CV^2^) is plotted against the mean expression level (normalized counts). Magenta points indicate mosquito genes showing higher than expected expression variability compared with ERCC spike-ins (blue) (adjusted *p* value <0.1). The red line is the fitted line of the spike-ins and the dashed line (pink) marks the margin for genes with 50% biological CV. (B) Pearson correlation heatmap of single hemocytes based on the expression of the highly variable genes identified in (A). Correlation suggests the presence of two groups of cells (red and yellow). (C) PCA plot based on the expression of highly variable genes. The first two principal components are shown, and each point represents one single hemocyte. Two clusters were identified and correspond to the subgroups in (B). *PPO6* expression, as log10 (normalized counts +1), is overlaid onto the PCA plot. (D) Violin plots of *PPOs* and *RFP* expression in the identified groups.

Considering the variable genes, we carried out hierarchical clustering based on pairwise Pearson correlation, which suggested the presence of at least two groups of mosquito blood cells in our samples (Figure 2B). Principal component analysis (PCA) also yielded two cell populations, supporting our clustering results (Figure 2C). Interestingly, the expression levels of *PPO6* showed high variability, and the overlay of *PPO6* expression onto the PCA plot suggested that the two clusters were largely characterized by low and high expression of *PPO6*. Differences in the *PPO6* expression levels have been previously described by immunofluorescence microscopy (Bryant and Michel, 2016) but have not been associated with cell types. Moreover, from the ten *PPO* genes encoded in the *A. gambiae* genome, eight were observed in our sequencing and six of them, as well as the *RFP* reporter, had variable expression between individual cells and the groups (Figure 2D and Table S3). Hence, we designated these two groups as PPO6^high^ and PPO6^low^.

### PPO6^high^ and PPO6^low^ cells represent transcriptionally distinct subpopulations

Among the highly variable genes, we detected several *FBN* sequences, such as *FBN8, 10* and *30*. PPO6^high^ cells showed high expression levels of *FBN10* (Figure 3A, left panel), whereas PPO6^low^ cells exhibited weak or lack expression of *FBN8, 10* and *30* (Figure 3A, Table S3). Although below the ERCC-defined variability threshold, likely due to the small number of cells analyzed, expression of the antimicrobial peptide gene *lysozyme type I (LysI)* (AGAP011119) was more characteristic of PPO6^low^ cells (Figure 3A, middle panel). In the search for a pan-hemocyte marker, we also identified expression of phagocytic receptor *Nimrod* in both groups of cells (Figure 3A, right panel). To validate the *in silico* data, we performed single-molecule RNA fluorescence *in situ* hybridization (RNA-FISH), and observed co-expression of *tdTomato* and *PPO6* in all *PPO6::RFP* hemocytes, with no detection of *tdTomato* in blood cells isolated from wild-type mosquitoes (Figure S2A-B, and data not shown). In terms of *PPO6* levels, RNA-FISH accurately distinguished PPO6^high^ and PPO6^low^ hemocytes, confirming our bioinformatics results. Consistently, PPO6^high^ cells showed high levels of *FBN10,* which were very low or absent in PPO6^low^ cells. High levels of *LysI* were found in PPO6^low^ cells, reinforcing the presence of PPO6^low^/FBN10^low^/LysI^high^ cells; and *Nimrod* was near ubiquitously expressed in all perfused hemocytes (Figure 3B). We took advantage of the high conservation levels of *PPO6, LysI* and *Nimrod* in the closely related mosquito species, *Anopheles stephensi,* the Asian malaria vector, to explore whether the newly discovered blood cell subgroups were present in other anopheline mosquitoes. Indeed, PPO6^high^ and PPO6^low^/LysI^high^ cells were seen, along with a low degree of expression of *Nimrod.* No *FBN10* was measured (Figure S2C), probably due to the specificity of the probe to *A. gambiae* and the large diversity of this gene family.

**Figure 3:**
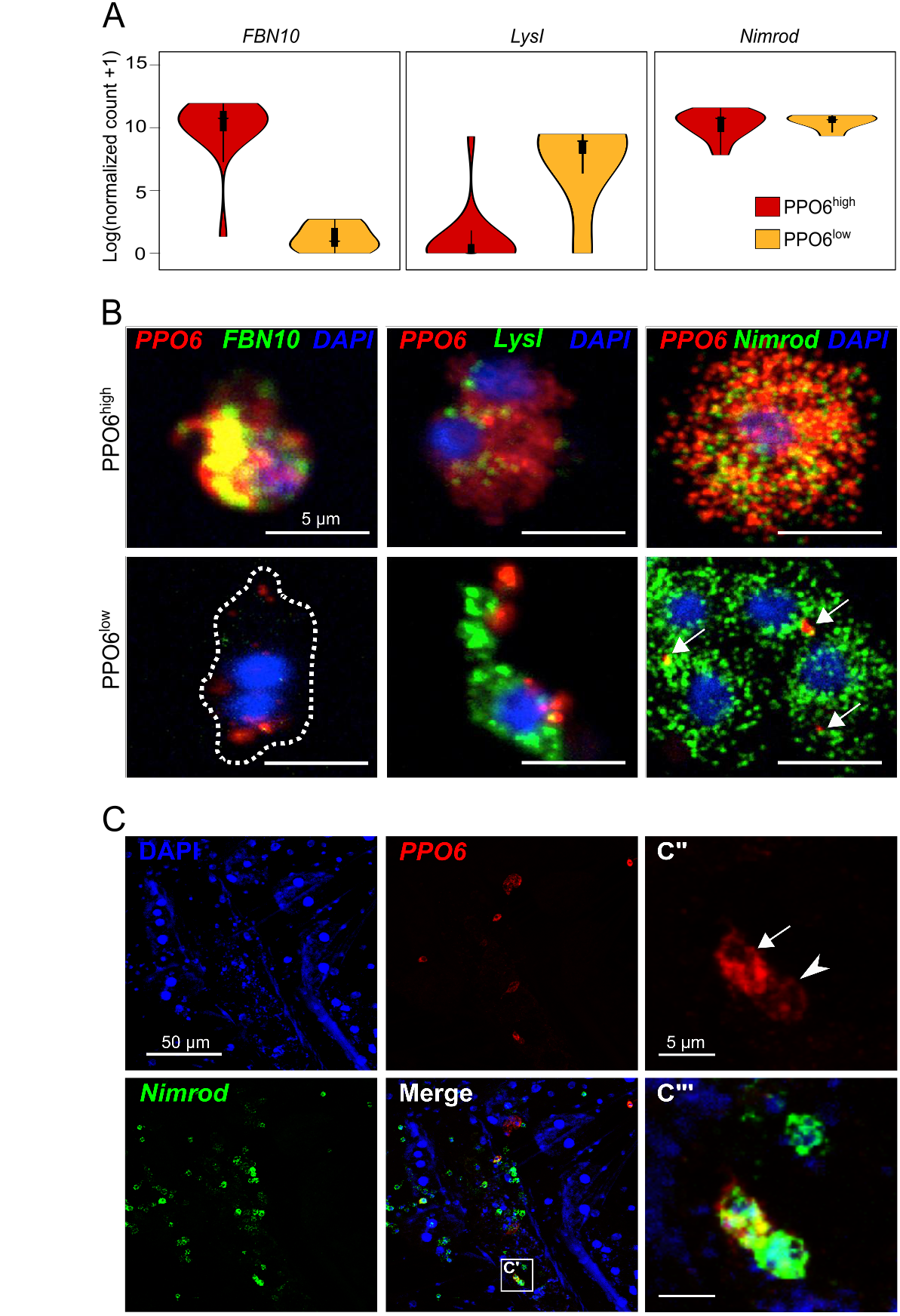
Characterization of PPO6^high^ and PPO6^low^ cell subpopulations. (A) Violin plots of the expression of putative population and pan-hemocyte markers. (B) RNA-FISH validation of identified PPO6^high^ and PPO6^low^ cell subpopulations in perfused cells based on markers shown in (A). Cells were classified as PPO6^high^ (upper panel) or PPO6^low^ (lower panel) according to the expression of *PPO6* (red). Arrows indicate lower *PPO6* signal. (C-C’’’) PPO6^high^ and PPO6^low^ cell subpopulations can also be seen as tissue-resident blood cells attached to the inner abdominal wall of female mosquitoes. Arrow and arrowhead indicate PPO6^high^ and PPO6^low^ cell subpopulations, respectively. Scale bars: (B) 5 μm, (C) 50 μm, (C’’-C’’’) 5 μm. DNA is stained with DAPI.

To obtain the transcriptional signatures of PPO6^high^ and PPO6^low^ cells, we compared the overall gene expression between these two groups. Based on differentially expressed genes, gene ontology (GO) analyses uncovered that melanization characterized PPO6^high^ cells, whereas metabolism and RNA processing defined the PPO6^low^ subset (Table S4). Although not significant, PPO6^low^ cells appeared to express more genes in total, but mitochondrial counts did not differ between the groups (Figure S2D). These findings suggest that PPO6^high^ cells are specialized for melanization responses, expressing genes involved in these processes at very high levels, whereas PPO6^low^ cells execute a broader range of biological tasks, including melanization. An alternative explanation would be that these groups represent different cell lineages, or perform melanization for diverse processes, e.g. metamorphosis and cuticle sclerotization, as established in other insects (Dudzic et al., 2015; Tsao et al., 2015; Tsao et al., 2009). Transcriptional differences between the two groups could also reflect localization patterns of the cells inside the mosquito body, as blood cells can be found in the circulation or attached to internal organs (King and Hillyer, 2013) and sessile cells could have been displaced during hemolymph perfusion. To assess whether observed differences were related to tissue residency, we performed RNA-FISH in tissues. The analyses of sessile cells revealed both cell populations in close contact with fat body cells within the abdominal wall with no conspicuous cell clusters. The majority of the sessile cells were positive for *Nimrod*, independent of their *PPO* expression (Figure 3C), indicating that *Nimrod* is a potential marker for both circulating and tissue-resident blood cells. Altogether, these results demonstrate that both circulatory and tissue-resident hemocytes display transcriptional heterogeneity, and that PPO6^high^ and PPO6^low^ cell populations are present in two mosquito species.

### PPO6^high^ and PPO6^low^ cells share functional and morphological features

Mosquito blood cells are functionally and morphologically separated into three classes - granulocytes, oenocytoids and prohemocytes - based on microscopic studies of surface-attached cells. Our GO analyses suggested the presence of a PPO-specialized cell population and a second cell subset of a less specific nature. Based on that, we reasoned that PPO6^high^ and PPO6^low^ cell groups could be representatives of oenocytoids and granulocytes, respectively. As phagocytosis is a hallmark of granulocytes, we first explored functional differences between these cells using magnetic bead uptake as means for “phagocyte” isolation, as suggested before (Smith et al., 2016). To this end, we injected mosquitoes with magnetic beads and either allowed them to rest at 28°C prior to hemolymph perfusion or incubated the mosquitoes at 4°C to inhibit phagocytosis. To our surprise, both PPO6^high^ and PPO6^low^ cells were identified among magnetically isolated cells and under both conditions, suggesting that instead of phagocytosis, both cell types endocytosed the beads, as no differences in bead uptake were observed when this assay was performed under cold conditions (Figure 4A, arrows). To confirm these findings, we compared the gene profiles of PPO-producing cells to the proteomics results obtained by Smith et al using magnetic bead isolation at 28°C only (Smith et al., 2016). Our analyses revealed that similarities were the strongest when profiles were compared across all samples - PPO6^high^, PPO6^low^, phagocytes and all cells, i.e. unselected cells obtained in the absence of magnetic isolation. PPO6^high^ and PPO6^low^ shared expression of more genes with phagocytes when considered together rather than alone, indicating that neither PPO6^high^ and PPO6^low^ cell types shared striking similarities with phagocytes at the gene/protein level (Figure 4B). In agreement, nearly all PPOs were present in all samples (Figure 4B’). Overall, these findings did not detect functional differences between PPO6^high^ and PPO6^low^ cells. We conclude that endocytic capabilities likely characterize both granulocytes, considered the “true” phagocytic cell type, and oenocytoids, known as the major source of PPOs.

**Figure 4:**
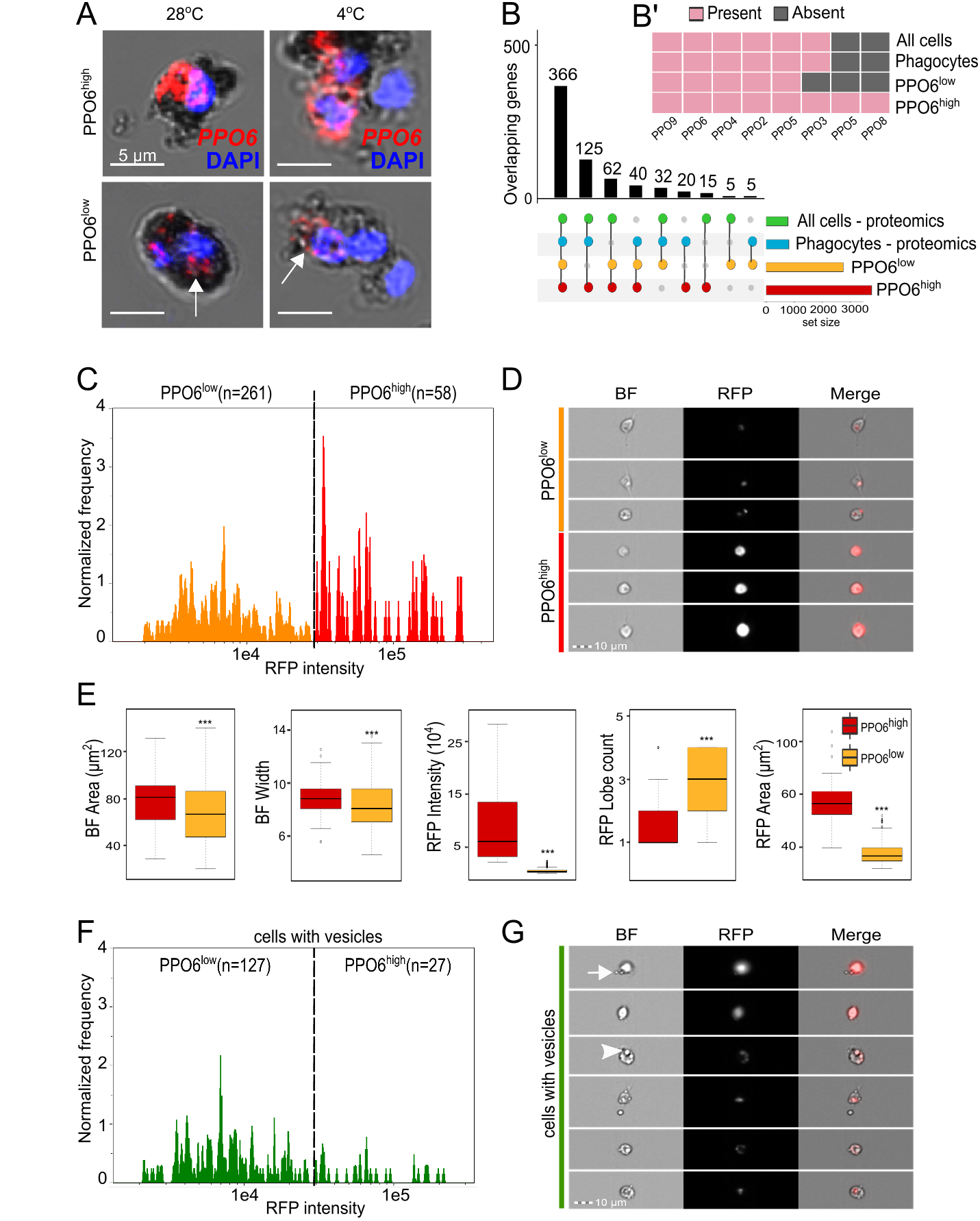
PPO6^high^ and PPO6^low^ cell subpopulations share functional and morphological features. (A) Magnetic bead isolation assays followed by RNA-FISH for identification of cell subpopulations. Arrows indicate lower *PPO6* signal. (B) Intersection analyses between PPO6^high^ and PPO6^low^ cell subpopulations and proteomics results obtained by Smith et al for phagocytes and all cells (unselected). Presence or absence of PPO genes/proteins in each of the samples in shown in (B’). (C) Hemocytes were perfused from *PPO6::RFP* mosquitoes and analyzed using imaging flow cytometry. RFP-positive cells were identified by their RFP Median Pixel and further separated into PPO6^high^ and PPO6^low^ based on their level of RFP fluorescence. The number of cells analyzed per group is shown between parentheses and the dotted line indicates the RFP threshold level used for separation of the two populations. (D) Image gallery containing representative images of PPO6^high^ and PPO6^low^ populations. Variation in cell shape and RFP intensity can be observed in the bright field (BF), RFP and merged images. (E) For every event measured within the flow, a corresponding image and fluorescence levels were acquired and analyzed to generate multiple morphological features. Boxplots show the distribution of five morphological measurements according to the groups. Asterisks represent p <0.01 based on Mann-Whitney-Wilcoxon test. (F) RFP-positive cells were interrogated for the presence of membrane protrusions or internal structures suggestive of vesicles. Similar to (C), cells were grouped into PPO6^high^ and PPO6^low^ and a RFP intensity histogram of cells displaying vesicles (green population) was generated to illustrate that vesicles are observed in cells from both groups. Representative images of cells in this population subset are shown in (G). Arrow and arrowhead indicate representative membrane protrusions and internal vesicles, respectively. See also Figure S4 and Tables S4-5. Boxplots indicate the median, first and third quartile, and min and max values.

Next, we sought to investigate whether the identified cell populations could be distinguished based on their morphology using imaging flow cytometry and RFP fluorescence as a proxy for *PPO6* expression. We measured a series of morphological features of 319 single RFP-positive cells, which could be divided into RFP^high^ (PPO6^high^, n=58) and RFP^low^ (PPO6^low^, n=261) based on their fluorescence intensity (Figures 4C-D and Figure S3). Overall, RFP-positive cells had a mean area of approximately 67 μm^2^, ranging from 18 μm^2^ to nearly 140 μm^2^. These measurements are in accordance with cell sizes reported elsewhere (Bryant and Michel, 2014, 2016; Castillo et al., 2006). Similar to recent studies based on flow cytometry of fixed cells (Bryant and Michel, 2014, 2016), we did not detect the cells of 2 μm in size described by other research groups based on label-free light microscopy alone (Rodrigues et al., 2010; Smith et al., 2015). When comparing the cell groups in terms of bright-field measurements of their cytoplasm (see Methods), PPO6^low^ cells showed smaller cytoplasmic area, width, and minor axis than PPO6^high^ cells (Figure 4E and Figure S3). No differences between the groups were detected in granularity or cell shape (Figure S8 and Table S5). To our surprise, PPO6^high^ and PPO6^low^ cells were equally circular. At times, cells from both groups also displayed an elongated shape, typical of the cytoplasmic extensions seen in fusiform or spindle-shaped cells. This fusiform shape is characteristic of plasmatocytes, described in other insects (Ribeiro and Brehelin, 2006). These results failed to morphologically assign RFP-positive cells to any of the morphologically-defined groups: granulocytes, plasmatocytes, or oenocytoids. In fact, the highest discriminating factors (Fisher’s linear discriminant, see Methods) separating PPO6^high^ and PPO6^low^ subpopulations relied on RFP intensity alone, with bright-field parameters scoring poorly and failing to establish a morphological distinction between the cells (Table S6). Importantly, our imaging flow cytometry approach relied on morphological analyses of cells in suspension, which is unbiased and likely more relevant for the identification of the cellular types found in the hemolymph circulation. This might explain differences obtained relative to previous studies that focused on cells attached to glass slides, which might be rather reflective of cell ‘states’ driven by activation of the cellular attachment and adhesion machineries upon exposure to electrostatically charged glass.

In addition to RFP intensity, the cell groups differed in their RFP area. PPO6^high^ cells displayed an overall cytoplasmic distribution of the RFP signal, whereas a more localized and globular signal was detected in the cytoplasm of PPO6^low^ cells, where RFP lobes were also more numerous, reinforcing the localized nature of the RFP signal (Figure 4D-E). Microscopic examination also revealed that nearly half of the cells from both groups displayed internal structures and or “budding” extensions of the cytoplasm suggestive of vesicles (Figure 4F-G, arrowhead and arrow, respectively). To confirm that, we performed correlative scanning electron microscopy (SEM) and demonstrated the presence of membrane protrusions or “blebs” in RFP-positive cells (Figure S4A). Altogether, these results established that morphological plasticity of the mosquito blood cells is independent from their transcriptional profile, and that mosquito blood cells have membrane vesicles and protrusions.

### Mosquito blood cells exchange molecular information

Mosquito hemocytes have been described to produce and secrete EVs (Castillo et al., 2017; Hillyer and Christensen, 2002) and, in agreement, our initial observation suggested that PPO-producing cells display vesicles. We were, therefore, puzzled by the possibility that the RFP signal analyzed in our cell sorting approach could have partially originated from RFP-positive vesicles. Earlier reports used DiD, a lipophilic cyanine dye, to label both mosquito hemocytes and hemocyte-derived vesicles (Castillo et al., 2017; King and Hillyer, 2012). To test whether the localized RFP signal seen in our imaging flow cytometry was associated with vesicles, we first stained PPO-producing cells with DiD and observed that RFP-positive cells indeed contained DiD-positive membrane-bound and internal vesicles that were both RFP-positive and negative (Figure S4B-C). Recently, it has also become clear that EVs are present in extracellular fluids like milk, saliva and plasma, so we reasoned that EVs might be also found in the hemolymph and in association with RFP-negative cells. To identify EVs in the mosquito circulation, we performed imaging flow cytometry analyses using DiD and a recently published approach (Headland et al., 2014). Both DiD-positive cells and EVs could be identified in hemolymph perfusate (Figure 5A-B). EVs were detected based on their small size, weak dark-field and positive DiD fluorescence, with a few EVs also displaying weak RFP signal (Figure 5B, arrowhead). The degree of DiD intensity differed between cells and did not depend on RFP fluorescence. Differential centrifugation followed by electron microscopy confirmed the presence of EVs in hemolymph perfusate of naïve female mosquitoes (Figure 5C). Moreover, SEM of perfused cells also revealed that vesicles of different sizes and shapes, suggestive of the different vesicle types described in the literature - exosomes, microvesicles and apoptotic vesicles (van der Pol et al., 2012), could be indeed observed in association with naïve mosquito blood cells (Figure S4D). These findings suggested that extracellular vesicle production is a general phenomenon that is not limited to PPO-associated cells.

**Figure 5:**
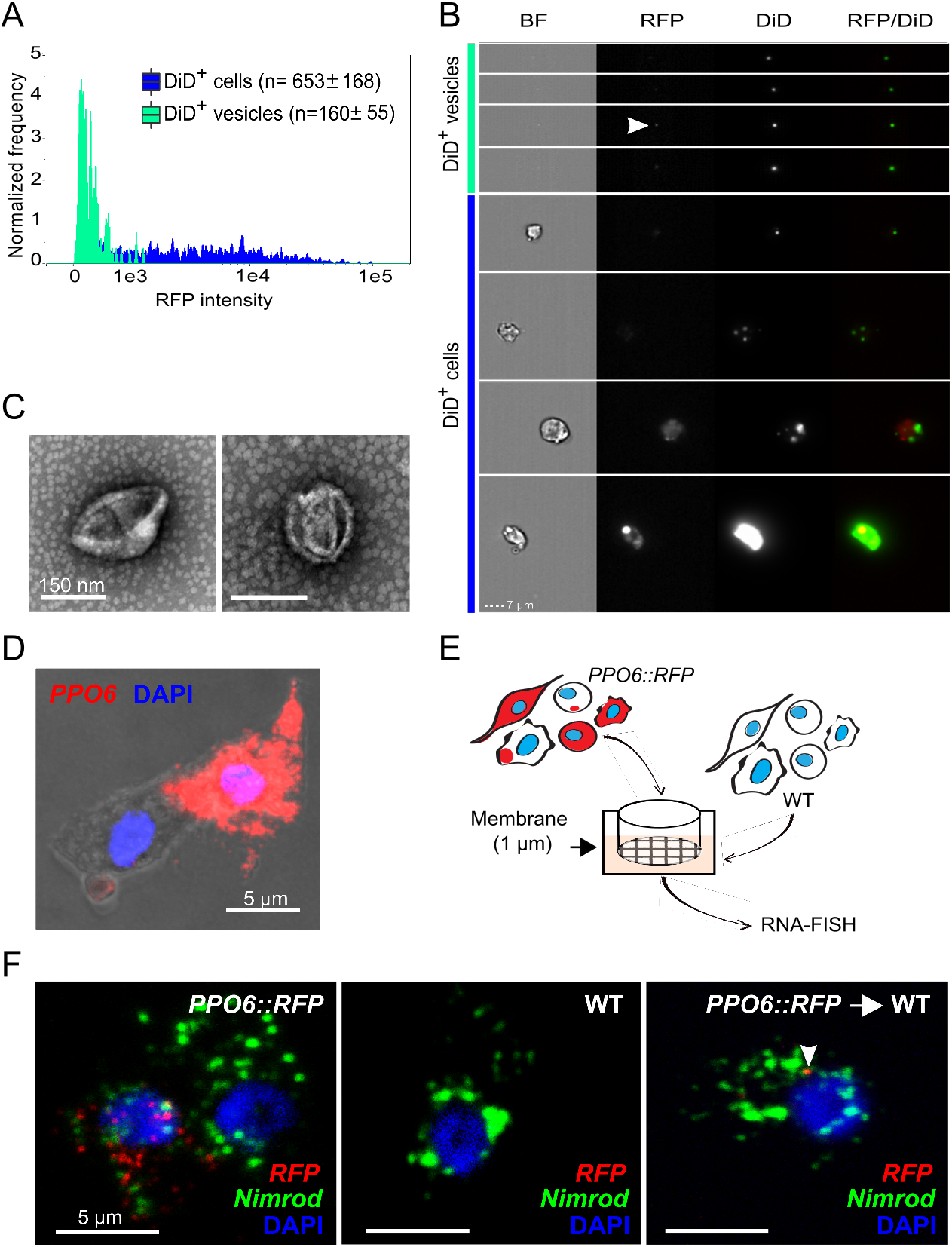
Vesicle identification and molecular exchange in mosquito blood cells. Hemocytes were perfused from *PPO6::RFP* mosquitoes, stained with the lipophilic DiD dye and analyzed using imaging flow cytometry. DiD-positive cells and EVs were first identified according to their DiD and darkfield intensity. EVs were further separated from cells based on their small bright field area. Cells were identified considering their area and aspect ratio. A RFP fluorescence histogram for both DiD-positive cells and EVs in shown in (A). (B) Representative images of DiD-positive cells and EVs. Arrowhead indicates a representative of a RFP-positive vesicle. (C) Negative staining electron microscopy of vesicles obtained by differential centrifugation (10,000xg) of hemolymph perfusate (representative images of 2 independent experiments are shown). (D) *PPO6* mRNA detection by RNA-FISH within budding extensions of blood cells. (E) Schematics of a transwell assay developed to test the transfer of *RFP* between blood cells from *PPO6::RFP* transgenic mosquitoes and wild-type mosquitoes that do not express any reporter genes. Hemolymph perfusate from wild-type mosquitoes was pipetted onto a coverslip placed under a 1 μm membrane. Perfusate collected from *PPO6::RFP* mosquitoes was placed on top of the membrane. (F) RNA-FISH was performed on coverslips obtained from the transwell assay described in (E) using probes to detect *RFP and Nimrod* expression in wild-type acceptor cells (right panel, arrowhead). Representative images of 2 independent experiments are shown. See also Figure S4.

A growing body of evidence has demonstrated that RNA can be transferred between mammalian cells. As RNA can be found in EVs, and our data showed that EVs are present in the mosquito hemolymph, we decided to take a step further and explore the possibility that a crosstalk between PPO-positive and negative cells could be responsible for the identification of PPO6^high^ and PPO6^low^ cells. Strengthening this idea were the observations that (1) our scRNA-seq results uncovered cells with minute levels of *PPO6* and *RFP,* and (2) expression of *PPO6* by RNA-FISH was detected inside “budding” extensions associated with PPO6-positive cells (Figure 5D). As a proof of concept, a transwell assay using blood cells from *PPO6::RFP* transgenic and wild-type mosquitoes was developed to test the possibility that RFP mRNA can be transferred between naïve transgenic and wild-type blood cells (Figure 5E). Remarkably, *RFP* transcripts were observed by RNA-FISH inside wild-type blood cells after exposure to hemolymph perfusate from transgenic mosquitoes (Figure 5F, arrowhead). This result indicated that *RFP* mRNAs can be shuttled between blood cells and might account for the PPO6^low^ cell population uncovered by our scRNA-seq and imaging approaches. Taken together, our findings demonstrated that molecular exchange between cells, likely via EVs, can affect their transcriptional profile. As EVs have been shown to carry lipids, proteins and RNA, and can be secreted by virtually all cells, our results revealed an unappreciated role of intercellular molecular exchange in defining cellular identity.

## DISCUSSION

Understanding how transcriptional networks influence cell identity is a central problem in modern molecular biology. Our study describes mosquito blood cells as a source of key components of immunity, development and tissue homeostasis, and places these cells as a central hub coordinating mosquito biology at different levels. Using a combination of single-cell genomics and imaging, we revealed that hemocytes display an unexpected degree of complexity where two transcriptionally defined cellular ‘populations’ suggestive of distinct cell types share morphological and functional features. We also demonstrate that mosquito blood cells exchange mRNA, leading to the detection, by RNA-FISH, of an “exogenous” gene in acceptor cells. Altogether, our results open a new perspective on cellular crosstalk and cell type classification, in addition to illustrating the power of single-cell-based approaches in discovering unappreciated events at the core of biological processes.

Using single-cell RNA sequencing, we describe the baseline expression of a mosquito blood cell in exceptional detail. An average mosquito blood cell under resting conditions expresses approximately 1,000 genes, or 7% of the mosquito transcriptome. In total, about half of the genes currently annotated in the mosquito genome were detected by RNA sequencing naïve, unstimulated mosquito hemocytes. This represents a substantial gene expression resource for further studies of tissue-specific alternative splicing, RNA editing, gene and transcript models. It will also enable the establishment of transcriptional and regulatory networks that allow for more precise gene enrichment and functional studies. In mosquitoes, where tissue-specific gene knockdown is not available and transgenesis remains limited, functional analyses have greatly relied on the use of dsRNA injections into the open body cavity. This technical limitation restricts the interpretation of tissue-specificity of signaling pathways and their regulation. We believe our dataset will illustrate the importance of tissue specificity studies and pave the road towards the detailed mapping of gene expression in cells and tissues of different systems, with the ultimate goal of creating a comprehensive reference atlas of cellular diversity.

By successfully applying single-cell RNA sequencing to the study of blood cells involved in immunity in a major malaria vector, we demonstrated proof of the existence of at least two transcriptionally distinct cell subpopulations that do not represent currently defined cell types. Our results show that the majority of *Anopheles* blood cells likely belong to the PPO6^low^ type. Their rich transcriptional program appears to be reminiscent of granulocytes. The second subpopulation, with a transcriptional profile specialized in melanization, is suggestive of oenocytoids. Nonetheless, these two transcriptionally defined cell types could not be distinguished by the customary functional or morphological tests. Therefore, we propose to call them plasmatocytes and melanocytes. Interestingly, only one cell type, plasmatocytes, is described in the adult *Drosophila* flies. Our transcriptional definition of mosquito blood cells redefines the obscure cellular classification extensively used in studies of hemocytes, and is particularly significant in the context of cellular proliferation and differentiation because mosquito hemocyte lineages are still to be established. Given our results, it will be interesting to address the spatial distribution of mosquito blood cells based on the markers identified here, especially as no lymph glands or hematopoietic cell clusters have been described in mosquitoes. Tissue-resident blood cells likely contribute to local responses and help regulate tissue-specific events. This is exemplified by the recent discovery of macrophage subsets regulating electrical pulsing in the mouse heart (Hulsmans et al., 2017), and of ovarian hemocytes that control collagen IV secretion and germline stem cell niche maintenance in *Drosophila* (Van De Bor et al., 2015). Evidence that cellular heterogeneity can be recognized by profiling single insect blood cells, often called macrophage-like, emphasizes the notion that innate immune cells are far more heterogeneous than previously thought. As macrophages, along with other immune cells, have their evolutionary roots in similar cells in ancestral invertebrates (Buchmann, 2014; Dzik, 2014) and new cell types arise as a result of evolutionary processes (Arendt et al., 2016), the study of insect blood cells can help elucidate the origins of the immune system.

In humans and mice, distinct levels and patterns of fluorescence of cellular markers are widely used in microscopy and flow cytometry as a means to cell type classification. Cellular crosstalk may, however, affect such approaches. Acquisition of macrophage-derived “blebs” by lymphocytes has been described, resulting in misrepresentation of lymphocytes as macrophages in flow cytometry studies, and suggesting that these two cells may interact to control early responses in the lymph node (Gray et al., 2012). More importantly, translation of transferred mRNAs into functional proteins has been demonstrated before (Valadi et al., 2007) along with the reprogramming of acceptor cells upon microvesicle-mediated exchange (Ratajczak et al., 2006). It is, therefore, intriguing to consider that expression of specific cellular markers might be influenced by EV uptake. In this scenario, signals sent via vesicles include protein-coding RNAs that are normally absent in acceptor cells, which once translated, appear to be an endogenous cellular marker characteristic of the donor cell population. This observation calls for a critical reassessment of cellular markers by the scientific community. It is also imperative to investigate whether certain protein-coding RNAs are preferentially exchanged compared to other RNAs found in the cells. How *PPO6* and *RFP* transcripts, and potentially other *PPO* genes, contribute to the function of acceptor cells is another exciting question. PPO proteins lack the signal peptides required for their secretion, and it has been suggested that PPO6 is secreted by exocytosis as cell rupture has not been observed (Bryant and Michel, 2016). It is plausible that PPO transcripts are shed by professional melanocytes and processed by non-professional plasmatocytes that locally activate melanization only under specific conditions, e.g. upon infection with specific pathogens, or during wounding and tissue repair. Molecular signals exchanged between cells can, thus, coordinate cellular plasticity and account for the diversity of functional subsets or ‘hybrid’ cells that express markers of different or multiple cell types.

The demonstration of mosquito blood-borne EVs indicates that different cells and tissues likely communicate through vesicles secreted into the insect open circulatory system. Several recent reports have suggested EV-mediated immune responses in dipteran insects. Exosome-like vesicles containing virus-derived siRNAs have been identified in *Drosophila* and contribute to systemic antiviral immunity (Tassetto et al., 2017). Apoptotic vesicles released by hemocytes in the proximity of invading parasites have been implicated in *anti-Plasmodium* responses by activating the complement pathway in *A. gambiae* mosquitoes (Castillo et al., 2017). Interestingly, using a GFP reporter strain, Volohonsky et al recently reported that, the anti-malaria mosquito complement-like factor TEP1 is predominately expressed in the fat body as a transcript, but at the protein level it is found in hemocytes upon blood feeding and infection (Volohonsky et al., 2017). The authors speculate this is due to the uptake by the blood cells of TEP1 attached to bacterial cells. As only one plasmatocyte containing low levels of *TEP1* was identified in our sequencing, we suggest that EV-mediated delivery of TEP1 (mRNA or protein) may better explain these surprising findings. We propose that vesicles found in the mosquito hemolymph contain proteins and transcripts that coordinate cell-to-cell and tissue communication not only in infection but also under physiological conditions. Disturbance in homeostasis, be it by infection, metabolic changes, tissue damage or stress, may further escalate secretion of vesicles containing an array of different cargo that can be targeted to specific tissues and complement systemic responses. This represents an outstanding parallel between insects and humans, where EVs found in human body fluids can elicit changes in function and gene expression in target cells, and have a direct role in processes like differentiation, neuronal signaling and cancer. We believe that, similar to how environment, microbiota and genetic make-up influence phenotypic variation, cellular exchange can also drive cellular identity and represents an inventive and unexplored way through which nature coordinates who and what we are.

## ACKNOWLEDGMENTS

The authors thank Suzana Zakoviz for sharing the drawing in Figure 1, and all members of the Vector Biology Unit for intellectual input and technical support. The authors are grateful to Dr. N. Regev-Rudzki for discussions and critical reading of the manuscript. The authors thank Dr. E. Marois (UPR9022 CNRS, U963 Inserm, France) for sharing the transgenic *PPO6::RFP* line, Dr. K. Müller (Humboldt University, Berlin) for providing *Anopheles stephensi* mosquitoes and acknowledge the discussions in the frame of the CNRS LIA “REL2 and resistance to malaria” project. We also acknowledge the support provided by the Flow Cytometry Core Facility (DRFZ/MPIIB, Berlin) and by Dr. I. Wagner (the Microarrays Core Facility, MPIIB, Berlin).

## AUTHOR CONTRIBUTIONS

M.S.S. and E.A.L. conceived the study, designed the experiments, analyzed the data and wrote the manuscript. M.S.S. performed all experiments with assistance from R.L.L. for the imaging flow cytometry studies, C.G. and V.B. for the EM analyses, and P.C. for the scRNA-seq. M.S.S. and J.J.M.L. performed bioinformatics analyses. J.H. developed web interface for data interrogation. A.E.H. provided infrastructure. V.B. and S.A.T. provided expert input.

## DECLARATION OF INTERESTS

The authors declare no competing interests.

## MATERIALS and METHODS

### Mosquito rearing, fluorescence microscopy and hemolymph perfusion

*Anopheles gambiae sensu lato PPO6::RFP* transgenic (Volohonsky et al., 2015) and wild type strains were reared at 28°C, under 80% humidity and at a 12/12 h day/night cycle. Larvae were fed with cat food and adult mosquitoes were fed *ad libitum* with 10% sugar solution. For tissue microscopy, adult female mosquitoes were dissected in 1 X PBS, fixed in 4% paraformaldehyde (PFA), washed three times and mounted using Vectashield mounting medium containing DAPI. For hemolymph perfusion, 3-5-day-old female mosquitoes were anesthetized on ice for 10 min, microinjected with 700 nl of a buffer containing 60% Schneider’s Medium, 10% fetal bovine serum (FBS), and 30% citrate buffer (anticoagulant; 98 mM NaOH, 186 mM NaCl, 1.7 mM EDTA, 41 mM citric acid, pH 4.5) and allowed to rest, on ice, for 10 min. A small cut was made between the last two abdominal segments with the help of dissection scissors and the flow through was collected after further injection of 10 μl of buffer. For microscopy analyses, mosquitoes were perfused directly onto glass slides or coverslips, and cells were allowed to attach for at least 15 min prior to fixation in 4% PFA. Cells were next stained with a 1:100 dilution of Alexa 488 Phalloidin (ThermoFisher) for 30 min at room temperature. For the DiD analyses, cells were stained with DiD (5 μM) for 20 min prior to PFA fixation. Following washes, cells were mounted as described above, and analyzed on a Zeiss Axiovert microscope.

### FACS and single-cell RNA-sequencing by SMART-Seq2

For FACS and imaging flow cytometry analyses, 10-12 mosquitoes were perfused. Hemolymph perfusate was collected with the help of a pipette, transferred into a siliconized microtube and diluted to a final volume of 500 μl buffer containing 2 μg/ml of Hoechst 3342 (Molecular Probes). Cells were immediately analyzed in a BD ARIA II Cell Sorter equipped with violet and green-yellow lasers at 405 and 561 nm, respectively. Cells were first gated based on their RFP fluorescence, followed by positive Hoechst signal, with Area versus Width measurements being used for doublet discrimination. The FACS machine was standardized with fluorochrome-containing beads and sorting purity was validated by visualization of cells sorted onto a glass coverslip. Cells were sorted into a 96-well PCR plate containing 5 μl of 0.2% Triton X-100 supplemented with 2 U/μl of RNAse inhibitor (Clontech), with two wells containing 30 cells each (pool samples) and one column (8 wells) containing only the lysis buffer used as a negative control. We added ERCC spike-ins (Ambion) at a 1:2 billion dilution into the plate prior to cDNA synthesis and all samples were next processed according to the SMART-Seq2 protocol using up to 22 PCR cycles for cDNA synthesis (Picelli et al., 2013). PCR products were purified with AMPure XP beads (Beckman Coulter). Quality control was performed for each sample individually both as cDNA input and sequencing library using a high sensitivity DNA kit (Agilent). A total of 125 pg of cDNA was used for library construction. cDNA libraries were pooled at a 10nM final concentration and 100 bp paired end sequencing was performed in one lane using a HiSeq2000 Sequencer (Illumina).

### RNA-seq data analysis

Reads generated by sequencing were demultiplexed using bcl2fastq (version 1.8.4) and mapped to *A. gambiae* genome (P4), ERCC92 (Ambion) and *dTomato* sequence (Volohonsky et al., 2015) with the STAR aligner (version 2.4.2a) (Dobin et al., 2013). The genome index was generated with *A. gambiae* geneset file in gtf format (P4.4) and gene count tables were produced during the mapping (–quantMode Genecounts). They were next normalized with size factors calculated from the ERCCs using DESeq2 (Love et al., 2014). For the purpose of comparisons, a gene was considered expressed if at least one normalized read was identified in at least one sample. Genes were annotated using Vectorbase (Lawson et al., 2007) and manual curation. For comparisons with previous studies (Baton et al., 2009; Pinto et al., 2009; Smith et al., 2016), gene IDs were converted using Vectorbase and BioMart. Intersection analyses were performed in R using the VennDiagram and upsetR packages. Technical noise estimation and identification of the highly variable genes were performed as reported before (Brennecke et al., 2013), using the 60- percentile as the mean cut-off to include more ERCC genes in the technical fit. Principal component analyses were done with the *prcomp* function and differential expression analyses were based on the DESeq2 package, using the ERCC size factors and PPO6^low^ versus PPO6^high^ as comparison. For GO analyzes, we used topGO (Alexa et al., 2006) and GO terms were obtained from the org.Ag.eg.db annotation package. Analyses were performed in R and scripts are available upon request. The sequencing results were deposited in the European Nucleotide Archive under the following accession number: PRJEB23372 and the processed expression data can be accessed at http://data.teichlab.org for single gene visualization.

### Imaging flow cytometry

Ten to twelve sugar-fed *PPO6::RFP* female mosquitoes were perfused to a final volume of 20-40 μl and the diluted hemolymph samples were immediately analyzed in an Amnis ImageStreamX MKII (Merck). For *PPO6::RFP* analyses, wild type mosquitoes were used to set background fluorescence and cells were measured with a 40x objective. Comparisons between cell subpopulations were performed using the “Object” mask and based on a built-in function that uses Fisher’s discriminant ratio (Rd) to determine the best statistical separation (largest Rd) between identified populations. For the DiD analyses, cells were collected into FBS-free buffer containing 1 μM of DiD and analyzed at 60x to increase resolution. Single staining controls representing RFP, DiD and medium alone were used for calibration and manual compensation. Experiments were repeated at least twice. Cell gating was confirmed considering the obtained images and manually curated to exclude debris and doublets that could not be excluded by the gating alone. Vesicle detection was performed as reported before (Headland et al., 2014). Briefly, DiD positive events were interrogated based on their level of DiD fluorescence and size scatter intensity. DiD-labelled vesicles showed a low scatter along with low to mid DiD fluorescence, whereas cells displayed mid to high fluorescence and scatter measurements. Speed beads, used for the instrument calibration and focusing, were easily gated out as a discrete population displaying very high levels of side-scatter intensity. The RFP intensity was measured based on the median pixel by means of histogram, and cellular debris and doublets were excluded according to their brightfield area and aspect ratio. The identified populations were compared using the “Feature Finder” function of the IDEAS software (MilliporeSigma). Statistical analyses were based on Mann-Whitney-Wilcoxon and graphs were done in R.

### RNA *in situ* hybridization using RNAscope

RNA *in situ* studies were performed according to the RNAscope Multiplex Fluorescence analyses manual (Advanced Cell Diagnostics). Briefly, cells were perfused onto glass slides, allowed to attach and fixed in PFA as described above. If needed, slides were dehydrated and kept in 100% ethanol at −20°C until processing. Samples from *A. stephensi* were handled following the same procedure. Tissue samples were processed immediately after dissection in RNAse-free PBS. All RNAscope probes were designed by Advanced Cell Diagnostics and are commercially available. Each probe was individually tested against a negative control prior and during each analysis. Images were acquired using a Leica SP8 confocal microscope equipped with 405, 488, 561 and 647 nm lasers and prepared for submission using the basic features of the LAS X software.

### Bead uptake assay

For magnetic isolation of hemocytes, we followed the protocol by Smith et al 2016 with small modifications. Briefly, 20 female mosquitoes were cold anesthetized and injected with 300 μl of a 2 mg/ml suspension of MagnaBind Carboxyl Derivatized Beads (Thermo Scientific). Mosquitoes were next kept at 28°C or 4°C for 2 h and perfused as described above. Hemolymph perfused was collected with the help of a pipette tip and transferred into a 0.5 μl Eppendorf tube containing 100 ul of injection buffer. Samples were diluted to 200 μl and incubated in a magnetic stand for 20 min at 4°C. Supernatant was removed by pipetting, magnetic pellet was resuspended in 1 X RNAse-free PBS and transferred to a microscopy slide. Cells were allowed to attach for 15 min and processed for RNA-FISH as described above.

### Transwell assay

For this assay, we used *PPO6::RFP* transgenic mosquitoes and the *A. gambiae* Ngousso (TEP1^*^S1) wild-type mosquito strain. Perfusion was performed in the absence of FBS and hemolymph was collected with the help of a pipette onto the top of a glass coverslip placed inside a 24-well plate. A total of 100 μl of buffer was further gently pipetted onto the cell drop to prevent dehydration. Cell inserts (Merck) were then placed over individual wells and hemolymph perfusate from *PPO6::RFP* female mosquitoes was gently pipetted onto the 1 μm membrane mesh. Diluted hemolymph from at least 2 wild type and 4 transgenic mosquitoes were used per treatment, and the negative control consisted of overlaying coverslips with buffer only. Plates were kept at room temperature for at least 1 h; following by fixation of coverslips with 4% PFA and PBS washes prior to immediate processing based on the RNAscope manual. Images were obtained by confocal microscopy as described above. Experiments were repeated at least twice.

### Scanning electron microscopy

For scanning electron microscopy (SEM), cells were perfused from at least 2 female mosquitoes directly onto glass coverslips and fixed with 4% PFA. To facilitate exosome imaging, poly-L-lysine treated coverslips were used. For correlative SEM, cells were placed onto microscopic dishes with finder grids (ibidi) and imaged directly after 4% PFA fixation using a Zeiss Axiovert microscope, prior to SEM processing. The samples were post-fixed in 2.5% glutaraldehyde, 0.5% osmium-tetroxide, tannic acid and osmium-tetroxide again. The coverslips or optical membranes were then dehydrated in a graded ethanol series, dried in carbon dioxide at critical point and vacuum coated with 3 nm Carbon-Platinum. Imaging was performed using a LEO 1550 (Zeiss, Oberkochen DE) scanning-electron microscope. Experiments were repeated at least twice.

### Transmission electron microscopy

EVs were isolated according to a protocol described before (Kowal et al., 2016). In summary, hemolymph perfusate from at least 20 mosquitoes was subjected to differential centrifugation at 4°C and the 10,000 × g pellet was processed for negative staining electron microscopy. Aliquots of the samples were applied to freshly glow discharged carbon- and pioloform-film-coated copper grids and allowed to adsorb for 10 min. After washes with distilled water, the grids were contrasted with 2% uranyl acetate, touched on filter paper and air-dried. The grids were examined in a LEO 906 (Zeiss AG, Oberkochen) electron microscope operated at 100 kV and images were recorded with a Morada (SIS-Olympus, Münster) digital camera.

